# Sequential regulatory activity prediction across chromosomes with convolutional neural networks

**DOI:** 10.1101/161851

**Authors:** David R. Kelley, Yakir A. Reshef, Maxwell Bileschi, David Belanger, Cory Y. McLean, Jasper Snoek

## Abstract

Models for predicting phenotypic outcomes from genotypes have important applications to understanding genomic function and improving human health. Here, we develop a machine-learning system to predict cell type-specific epigenetic and transcriptional profiles in large mammalian genomes from DNA sequence alone. Using convolutional neural networks, this system identifies promoters and distal regulatory elements and synthesizes their content to make effective gene expression predictions. We show that model predictions for the influence of genomic variants on gene expression align well to causal variants underlying eQTLs in human populations and can be useful for generating mechanistic hypotheses to enable fine mapping of disease loci.

## Introduction

Although many studies show strong relationships between variation in genotype and phenotype across a range of human diseases and traits, the mechanisms through which this relationship operates remain incompletely understood (**Boyle et al. 2017**). Noncoding variation has especially stifled progress; most genomic loci statistically associated with phenotypes via genome-wide association studies (GWAS) do not alter coding sequence, but mechanisms for only a rare few have been thoroughly described (e.g. (**Claussnitzer et al. 2015**)). Numerous lines of evidence suggest that many noncoding variants influence traits by changing gene expression (**Finucane et al. 2015; O’Connor et al. 2017; Albert and Kruglyak 2015; Maurano et al. 2012**). In turn, gene expression determines the diversity of cell types and states in multi-cellular organisms (**Roadmap Epigenomics Consortium et al. 2015**). Thus, gene expression offers a tractable intermediate phenotype for which improved modeling would have great value.

Large-scale consortia and many individual labs have mapped the epigenetic and expression profiles of a wide variety of cells (**The ENCODE Project Consortium et al. 2012; Roadmap Epigenomics Consortium et al. 2015; Forrest et al. 2014**). Further, it has recently become appreciated that many of these data can be accurately modeled as a function of underlying DNA sequence using machine learning. Successful predictive modeling of transcription factor (TF) binding, accessible chromatin, and histone modifications has provided mechanistic insight and useful interpretation of genomic variants (**Whitaker et al. 2015; Ghandi et al. 2014; Alipanahi et al. 2015; Zhou and Troyanskaya 2015; Kelley et al. 2016**). In particular, the expansive training data available from the 3 billion-nucleotide human genome has enabled deep learning methods with huge numbers of parameters to make significantly more accurate predictions on held out test data than previous approaches (**Zhou and Troyanskaya 2015; Kelley et al. 2016**).

Despite this progress, models to predict cell type-specific gene expression from DNA sequence have remained elusive in complex organisms. Existing models use experimental annotations as input (e.g. peak calls for various known regulatory attributes), allowing them to shed light on the relationships between these annotations (**González et al. 2015; Cheng et al. 2012**), but not analyzing the causal role of the underlying sequence in determining those annotations. Even with intra-experiment training data to infer the relevant sequence motifs, the complexity of distal regulation where enhancers can interact with promoters across hundreds of thousands of nucleotides challenges the current generation of methods (**Long et al. 2016; Levine 2010**). However, well-established gene regulation principles from inquiry into enhancer biology and 3D chromosomal contact domains has yet to be fully incorporated into expressive machine learning models (**Dekker and Mirny 2016; Mifsud et al. 2015**). Modeling larger sequences and diverse experimental data offers a path forward to improve predictive accuracy. More effective models would enable researchers to profile one instance of a tissue or cell type and project that profile to individuals with varying genomic sequence.

Here, we use novel machine-learning algorithms to learn to predict thousands of epigenetic and transcriptional profiles from hundreds of human cell types using only DNA sequence as input. Using the model, we predict the difference between the two alleles of genomic variants for these thousands of datasets, focusing particularly on predicted changes to gene expression. We demonstrate the considerable potential value of this observation to identify likely causal variants and mechanisms within GWAS loci.

## Results

### Basenji

In previous work, we introduced a deep convolutional neural network approach named Basset for modeling “peak”-based chromatin profiles, focusing particularly on DNase I hypersensitivity (**Kelley et al. 2016**). Given an input sequence of 500-1000 bp, the model makes a single binary prediction for the sequence’s activity in any training dataset provided. Here, we modify the Basset architecture to (1) model distal regulatory interactions and (2) predict finer resolution, quantitative (as opposed to binary) genomic profiles that are more appropriate for the dynamic range of gene expression (Figure 1). As a related approach, but one begging for metaphor to a more nimble and far-sighted hound, we refer to this method as Basenji.

**Figure 1.**
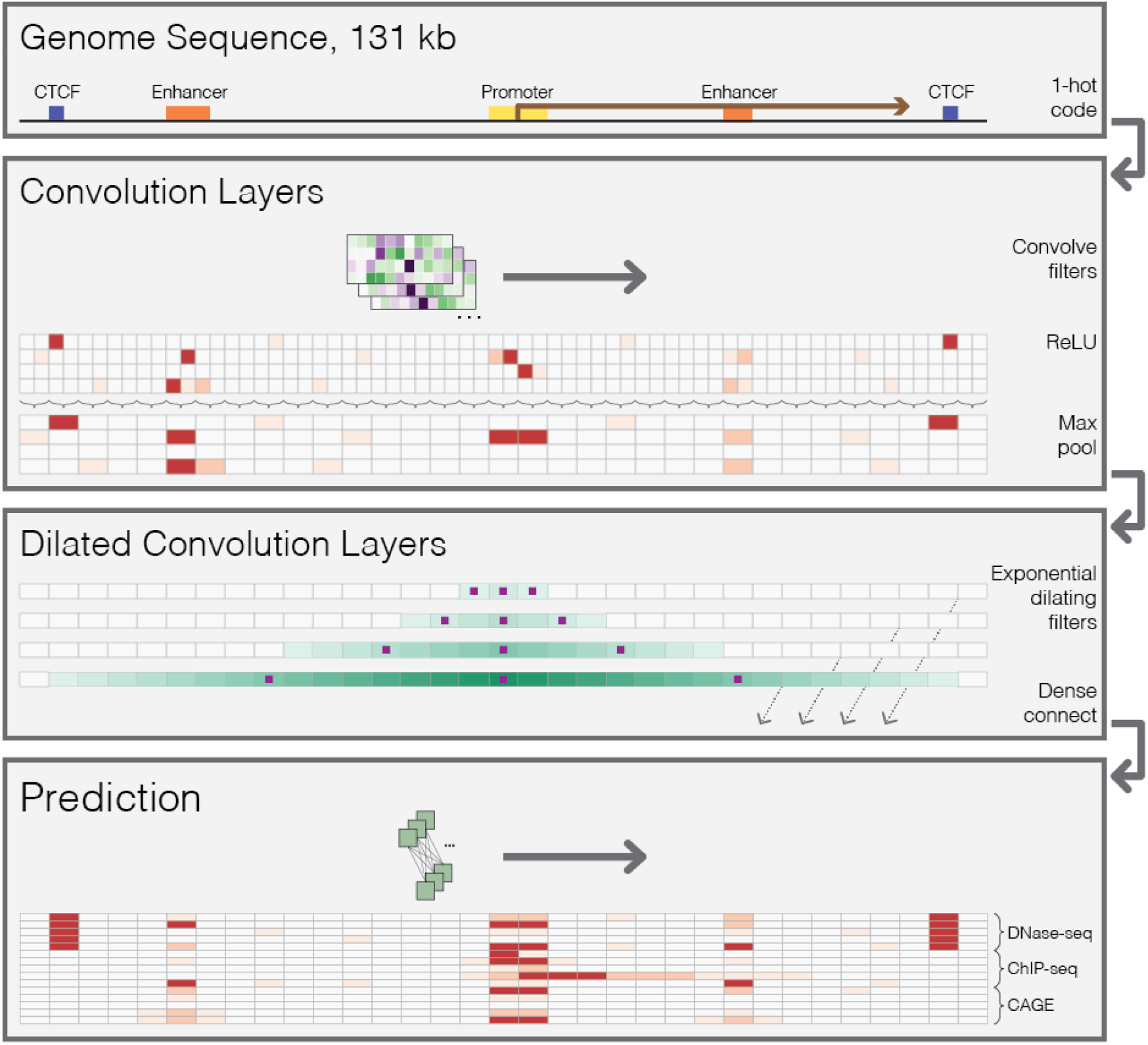
Sequential regulatory activity prediction. DNA sequences come in to the model one hot coded to four rows representing A, C, G, and T. The annotations are fabrications to help convey the reasons for the various elements of the architecture. We apply several layers of convolution and max pooling, similarly to previous methods (Kelley et al. 2016), to obtain representations that describe 128 bp bins. To share information across large distances, we apply several layers of dilated convolutions. The purple squares indicate the columns that the convolution directly sees; the teal shade is drawn proportional to the number of operations performed on that column with respect to the center position. Dilated convolution layers are densely passed on to the final prediction layer, where a width-one convolution layer makes predictions across the sequence. We compare these predictions to the experimental counts via a Poisson regression loss function and use stochastic gradient descent with back propagation to fit the model parameters.

The model accepts much larger (2^17^=)131 kb regions as input and, similarly to Basset, performs multiple layers of convolution and pooling, to transform the DNA to a sequence of vectors representing 128 bp regions (Methods). To share information across long distances, we then apply several layers of densely connected dilated convolutions (Methods). After these layers, each 128 bp region aligns to a vector that considers the relevant regulatory elements across a large span of sequence. Finally, we apply a final width-one convolutional layer to parameterize a multi-task Poisson regression on normalized counts of aligned reads to that region for every dataset provided (**Hashimoto et al. 2016**). That is, the model’s ultimate goal is to predict read coverage in 128 bp bins across long chromosome sequences.

Modeling count data, as opposed to peak data, required careful preprocessing beyond that performed in the standard pipelines of genomics consortium projects. For example, processed consortium data discards multi-mapping read alignments, which leaves half the genome incompletely annotated, despite the substantial evidence that repetitive sequence is critical to gene regulation (**Feschotte 2008**). Thus, we downloaded the original sequencing reads for 593 ENCODE DNase-seq, 1704 ENCODE histone modification ChIP-seq, 356 Roadmap DNase-seq, 603 Roadmap histone modification ChIP-seq, and 973 FANTOM5 CAGE experiments. These experiments represent 529 unique cells/tissues profiled by DNase-seq, 1136 unique cells/tissues profiled by ChIP-seq, and 595 unique cells/tissues profiled by CAGE. We processed these data with a pipeline that includes additional computation to make use of multi-mapping reads and normalize for GC bias (Methods). Though additional data modalities may require slight modification, this base pipeline will allow seamless addition of future data.

We trained to fit the neural network parameters on one set of genomic sequences annotated by all datasets and benchmarked predictions on those same datasets for a held-out set of test sequences. We performed all analyses in this manuscript on the test sequences. We used a Basenji architecture with 4 standard convolution layers, pooling in between layers by 2, 4, 4, and 4 to a multiplicative total of 128, 7 dilated convolution layers, and a final convolution layer to predict the 4229 coverage datasets. We optimized all additional hyper-parameters using Bayesian optimization (Methods).

### Prediction accuracy

To assess how effectively the model predicts the signal in these datasets, we compared predictions to experimental coverage in the 128 bp bins for a set of held-out test sequences (Figure 2A). As previously observed, accuracy varies by the type of data—punctate peak data tend to be more directly dependent on the underlying sequence, making for an easier prediction task (Figure 2B, 2C) (**Whitaker et al. 2015; Zhou and Troyanskaya 2015**). Accordingly, Basenji predictions explained the most variance in DNase-seq and ChIP-seq for active regulatory regions. Lower accuracy for the broad chromatin domains marked by modifications like H3K79me2 and H3K9me3 is expected because they depend more on distant sequence signals and incompletely understood propagation mechanisms (**Hathaway et al. 2012**).

**Figure 2.**
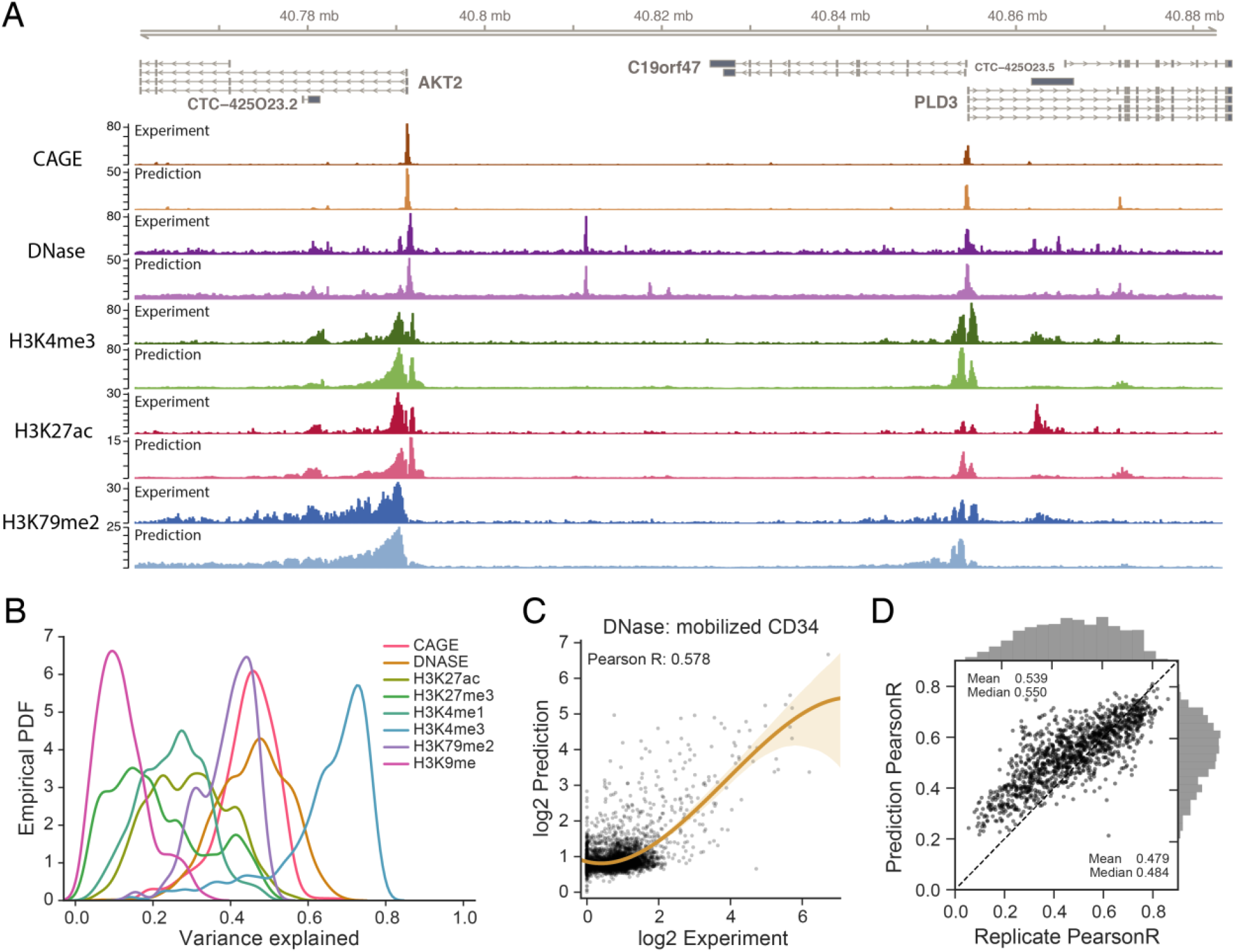
Basenji predicts diverse epigenetic and transcriptional profiles from DNA sequence. (A) The AKT2 locus exemplifies the genome-wide accuracy of Basenji predictions; gene promoters and the strongest distal regulatory elements are easily identified, with some false positive and negative predictions for weaker elements. For each track, the darker version on top represents the experimental coverage and the lighter version below represents Basenji predictions. (B) We computed the variance explained (R^2^) for each experiment and plot here the distributions classified by dataset type. Basenji predicts punctate peak data, but broad chromatin marks remain challenging. (C) For the median accuracy DNase-seq experiment, mobilized CD34 cells, we plotted the log_2_ predictions versus log_2_ experiment coverage in 128 bp bins. (D) For all replicated experiments, we plotted log-log Pearson correlation between the replicate experiments versus the correlation between the experiment and its replicate’s prediction (averaged across replicates). Both the mean and median Basenji prediction accuracy exceed the replicate accuracy.

To benchmark this framework against our previous Basset (**Kelley et al. 2016**), we focused on the 949 DNase-seq experiments, divided the longer sequences into 1024 bp subsequences, and classified each subsequence in each experiment with a peak calling algorithm (Methods). We trained a Basset model using a set of hyper-parameters studied in a recent application of the method (**Reshef et al. 2017**). Basenji quantitative predictions achieved a greater AUPRC for all experiments, increasing the average from 0.435 to 0.577 and median from 0.449 to 0.591 (Supplemental Fig S1).

1284 replicated experiments allowed us to benchmark prediction-experiment correlation versus experiment-experiment correlation. Determining the exact nature of these replicates was not always possible, but 80-90% are biological (rather than technical) replicates. Across these replicated experiments, the mean and median prediction correlation exceeded the replicate correlation (paired t-test p-value < 2×10^−78^) (Supplemental Fig S2). The mean and median cross-replicate predictions (i.e. the prediction for replicate one versus the experimental data for replicate two) also exceeded the correlation between replicates (paired t-test p-value <7×10^−7^) (Figure 2D; Supplemental Fig S3). As replicate-replicate correlation increased, Basenji prediction accuracy also increased, suggesting that higher quality data enables more effective modeling of the sequence dependence of the regulatory signal (Figure 2D). The silencing modifications H3K9me3 and H3K27me3 had low replicate consistency, and improved data may lead to better modeling of repressive chromatin in the future (Supplemental Fig S2,S3). Given the limited focus on transcriptional regulation, Basenji prediction correlations approach but fall short of the replicate-replicate correlations at the high end for the most consistent datasets, particularly CAGE (Figure 2D; Supplemental Fig S2,S3).

We hypothesized that the 7 dilated convolution layers enabled the model to capture the distal influences that are an established feature of human gene regulation. To isolate the influence of receptive field width, we trained similar models with 1-7 dilated layers. Test accuracy increased with increasing receptive field for all data types, confirming the value added by the additional dilated convolution layers of the final network (Supplemental Fig S4). An 8^th^ layer would reach outside the bounds of the sequence too frequently for this input length, but may add value for longer sequences.

### Cell type-specific gene expression

A driving goal of this research is to effectively model cell type-specific gene expression. CAGE quantifies gene expression by capturing and sequencing 5’ capped mRNAs to measure activity from genes’ various start sites (**Forrest et al. 2014**). To offer a gene-centric view from Basenji predictions, we focused on annotated TSSs (GENCODE v25) and considered the experimental measurement and prediction in the 128 bp bin containing each TSS. For each gene outside the training set, we summed their various TSS values to compute accuracy statistics. After log_2_ transform, the mean Pearson correlation of gene predictions with experimental measurements across cell types was 0.85 overall and 0.77 for nonzero genes (Figure 3A,B) (medians 0.86 and 0.78 respectively), which is on par with models that consider measurements of the relevant regulatory events with sequencing assays rather than predicting them from sequence (**Cheng et al. 2012**). Correlation is greater for CAGE datasets with more reads aligned to TSSs (Figure 3A), suggesting that low CAGE sequencing depth constrains accuracy for many datasets.

**Figure 3.**
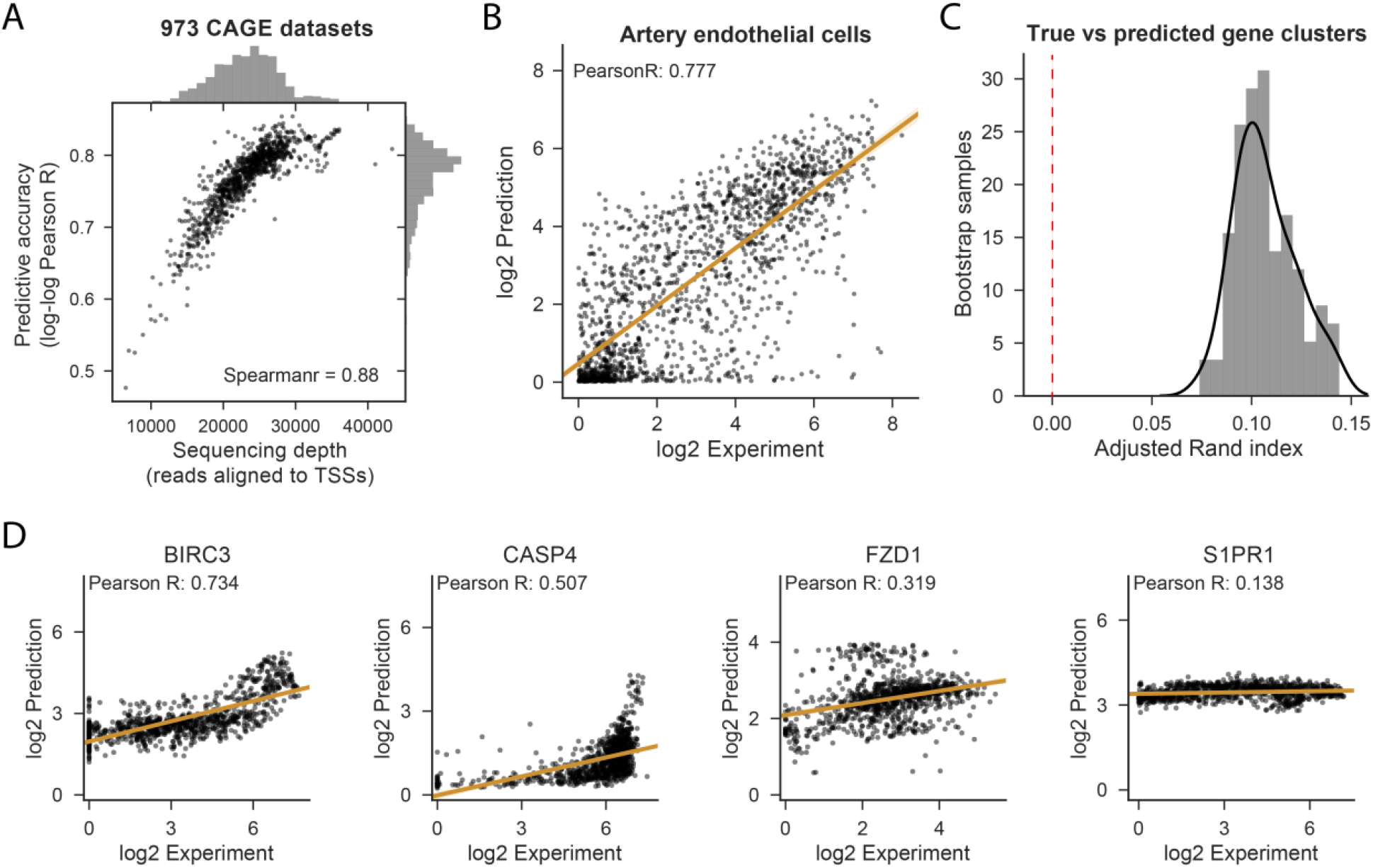
Basenji predicts cell type-specific gene expression. (A) We computed Pearson correlation between the log_2_ prediction and experiment across all nonzero expressed test set genes for each CAGE dataset. We plotted those correlations against the total number of reads aligned to test gene TSSs, which measures the relevant sequencing depth. (B) For the median accuracy cell, artery endothelial cells, we plotted the experiment coverage versus Basenji prediction. (C) For both the experimental measurement and Basenji prediction, the gene expression by CAGE dataset matrix displays clusters. We measured the similarity of those clusters between the experimental and predicted data by bootstrap sampling gene subsets, clustering both the experimental and predicted data, and computing the adjusted Rand index between the cluster sets (Methods). The adjusted Rand index is significantly greater than the null model value zero (p-value < 1×10^−26^). (D) We plotted gene expression versus prediction after quantile normalization across cell types for the genes ranked in the 95th, 75th, 50th, and 25th percentiles by Pearson correlation.

Predictions varied across cell types, suggesting that the model learns cell type-specific transcriptional programs (Supplemental Fig S5). For these analyses, we performed quantile normalization on the predictions and experimental measurements across datasets. To quantify concordance between cell/tissue cluster structure, we performed Gaussian mixture model clustering of bootstrap gene samples for the experimental and predicted expression profiles. The adjusted Rand index distribution (mean 0.107, median 0.104) indicated significant agreement between clusters (p-value < 1×10^−18^) (Figure 3C).

To quantify the greater difficulty of predicting highly cell type-specific expression, we computed Pearson correlation for sets of genes binned into quartiles by their coefficient of variation across all CAGE samples (Supplemental Fig S6). The stable expression across cell types of housekeeping genes relies on a particular promoter architecture (**Lenhard et al. 2012**), which the model learns well. Predictions for genes with medium levels of variability across samples have similar correlation levels. Only in the most variable quartile do we observe a deterioration in correlation from mean 0.3673 (median 0.3592) across targets in the third quartile to mean 0.2708 (median 0.2562) in the fourth. Nevertheless, the predictions explain considerable variance for even these most variable genes, suggesting that the model learns some degree of cell type specificity.

We found it instructive to closely examine genes with variable expression patterns. We computed accuracy statistics independently for each gene on their vectors of quantile normalized predictions and experimental measurements across cell types. In Figure 3D, we display genes at the 95th, 75th, 50th, and 25th percentiles ranked by correlation. Instances of effective predictions across several orders of magnitude, such as for *BIRC3* with its greatest expression in the small intestine, stomach, and spleen, lend credence to the model. In many cases, Basenji has learned that the gene’s expression varies across cells, but underestimates the dynamic range of the variance. For example, *CASP4* and *FZD1* predictions correlate with the experimental measurement, but compress the range between the most and least expressed cells. Some degree of variance reduction is warranted because the CAGE measurement includes stochastic noise that Basenji will implicitly smooth out. However, poor prediction of some genes, such as *S1PR1* indicate that an inability to capture more complex regulation also dampens prediction confidence.

CAGE quantifies gene expression at TSS resolution, and genes may use distinct TSSs in different cell types (**Ayoubi and Van De Ven 1996**). To measure Basenji’s ability to model these TSS switches, we delineated a set of 201 genes with TSSs >500 bp apart and variance across all CAGE samples >1 (using log_2_-transformed, quantile normalized expression estimates). For each gene’s two most variable TSSs, we computed the log_2_ fold change of one TSS versus the other in each CAGE sample for both the experiment and Basenji prediction. The mean Pearson correlation between experiment and prediction fold changes was 0.295 (median 0.288) over the set of genes (Supplemental Fig S7), which is greater than zero with p-value < 1×10^−90^ by *t*-test. This analysis suggests that Basenji predictions explain considerable variance in alternative TSS usage across samples.

### Distal regulatory elements

Distant enhancer sequences play a significant role in activating gene expression (**Long et al. 2016**). We devised a method to quantify how distal sequence influences a Basenji model’s predictions and applied it to produce saliency maps for gene regions. The saliency map scores annotate 128 bp segments with a function of the model predictions’ gradient with respect to that segment’s vector representation after the convolutional layers and before the dilated convolutions share information across wider distances (Methods). Peaks in this saliency score detect distal regulatory elements, and its sign indicates enhancing (+) versus silencing (−) influence.

The region surrounding *PIM1* in the GM12878 cell line exemplifies this approach (Figure 4A). The promoter region has extreme saliency scores, including repressive segments; i.e. mutating the regulatory sequence recognized by the model in these regions would increase the predicted activity. Elements beyond the promoter are also captured—we identified two enhancers annotated by ENCODE, one with a panoply of motifs highlighted by POU2F and another with two adjacent PU.1 motifs. We searched for perturbation data to support these regulatory interactions. In a collection of 59 siRNA TF knockdowns performed in a similar lymphoblastoid cell line GM19238, POU2F1 and POU2F2 knockdowns resulted in differential expression of *PIM1* mRNA with p-values 0.026 and 0.066 respectively (as tested by a likelihood ratio test within the framework of a fixed-effect linear model of batch-corrected microarray measurements; detailed Methods in (**Cusanovich et al. 2014**)).

**Figure 4.**
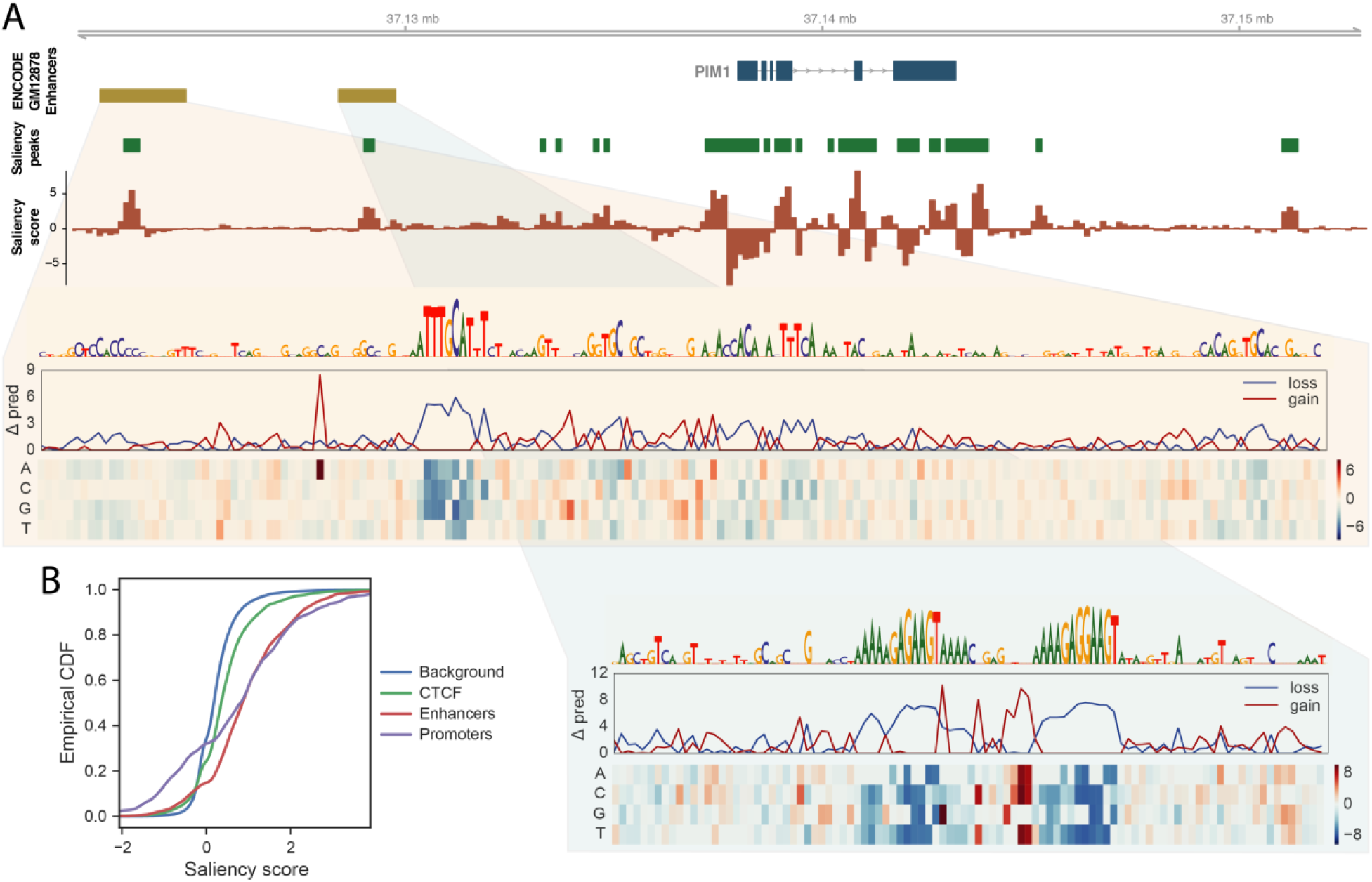
Basenji identifies distal regulatory elements. (A) ENCODE enhancer annotations for *PIM1* in GM12878 specify two downstream regulatory elements. Basenji saliency scores and FDR<0.05 peaks (see Methods) mark these elements, in addition to a variety of others that lack typical enhancer chromatin. In silico saturation mutagenesis of these elements with respect to Basenji’s *PIM1* GM12878 CAGE prediction outline the driving motifs. The quantities in the heatmap display the change in Basenji prediction “Δ pred” (summed across the sequence) after substituting the row’s specified nucleotide into the sequence. The line plots display the min (loss) and max (gain) change among the possible substitutions. The upstream cis-regulatory module most prominently features a POU2F factor motif, while the downstream element consists solely of two adjacent PU.1 motifs. (B) We plotted the cumulative distributions of the maximum saliency score for elements of various regulatory annotation classes in GM12878 released by ENCODE. Genome-wide, each annotation class differs significantly from the background scores by Kolmogorov-Smirnov test.

To assess this method’s ability to detect such regulatory elements genome-wide, we downloaded several curated annotations from ENCODE for GM12878—promoters, enhancers (not overlapping promoters), and CTCF binding sites (not overlapping promoters or enhancers) (**The ENCODE Project Consortium et al. 2012**). We computed the maximum saliency score overlapping instances of these annotations and shuffled background sets. Scores for promoters, enhancers, and CTCF sites differed significantly from scores for the background set (Kolmogorv-Smirnov test p-values 9×10^−64^, 2×10^−183^, and 4×10^−61^ respectively) (Figure 4B) (Methods). Intriguingly, promoters have more extreme scores at both the high and low ends, suggesting that they frequently contain repressive segments that may serve to tune the gene’s expression rate. This feature is also present for enhancers, but at a far lesser magnitude on the repressive end.

### Expression QTLs

Functional profiling of genotyped individuals is widely used to detect influential genomic variation in populations. The Gene-Tissue Expression (GTEx) project offers one such data collection (The GTEx Consortium 2017). GTEx measured RNA abundance via RNA-seq in 44 separate human tissues post-mortem and computed association tests for genomic variants significantly correlated with gene expression (eQTLs). Without observing such data, a trained Basenji model can be used to predict which single nucleotide polymorphisms (SNPs) are eQTLs by comparing model output for the different SNP alleles. To benchmark this approach, we downloaded the GTEx V6p release and focused on 19 tissues that were reasonable semantic matches for FANTOM5 CAGE profiles.

Given a SNP-gene pair, we define its SNP expression difference (SED) score as the difference between the predicted CAGE coverage at that gene’s TSS (or summed across multiple alternative TSS) for the two alleles (Figure 5A). Linkage disequilibrium (LD) complicates the comparison to eQTL statistics; marginal associations and significance calls depend on correlated variants in addition to the measured variant, and association scans are better powered for variants that tag more genetic variation (**Bulik-Sullivan et al. 2015**). To put SED on a level plane with the eQTL statistics, we distributed the SED scores according to variant correlations to form a signed LD profile of our SED scores, here denoted SED-LD (Methods) (**Reshef et al. 2017**).

**Figure 5.**
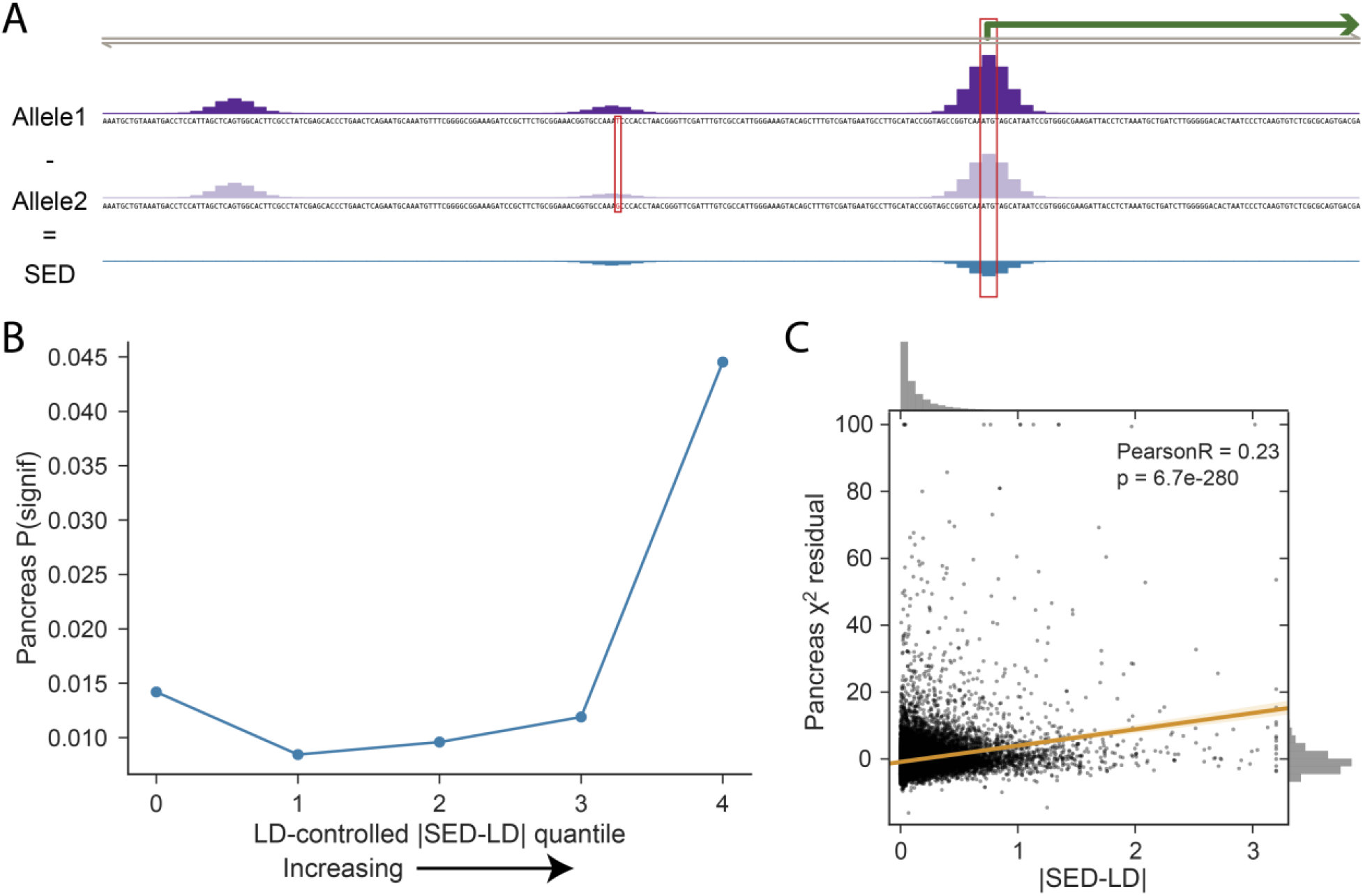
Basenji gene-specific variant scores enrich for eQTLs. (A) We defined SNP expression difference (SED) scores for each bi-allelic variant and gene combination as the difference between the model prediction for the two alleles at that gene’s TSSs. (B) We computed the signed LD profile of the SED annotations (denoted by SED-LD) to more readily compare to eQTL measurements in human populations (Methods). |SED-LD| shows a strong relationship with eQTL statistics from GTEx. Here, we binned variants into five quantiles by the difference between their regression predictions including and excluding |SED-LD| and plotted the proportion of variants called significant eQTLs in pancreas. The proportion rises with greater |SED-LD| to 4.2x in the highest quantile over the average of the bottom three quantiles, which represented the median enrichment in a range of 3.2-5.8x across the 19 tissues. See Supplemental Fig S8 for all tissues and TSS-controlled analysis. (C) Plotting |SED-LD| versus the chi-squared statistics reveals a highly significant correlation.

We checked whether the absolute value of SED-LD correlated with eQTL chi-squared statistics after controlling for the total amount of genetic variation tagged by each SNP as measured by LD score (**Bulik-Sullivan et al. 2015**) on a set of LD-pruned variants (Methods). Indeed, |SED-LD| significantly correlated with the adjusted eQTL statistics in all 19 tissues (p-values all <1×10^−54^ using LD-pruned variants from chromosome 1). To assess the quantitative extent of this enrichment, we ranked variants by the difference between their predictions from regression models including and excluding |SED-LD| for each tissue and binned into five quantiles. The proportion of variants called significant eQTLs was 3.2-5.8x greater in the top quantile relative to the average of the bottom three in all tissues (Figure 5B, 5C). This effect was robust to controlling for distance to TSS (Supplemental Fig S8). Thus, our analyses support a robust predictive relationship between Basenji scores and population measurements of RNA abundance, despite the additional layers of post-transcriptional regulation captured by the eQTL analysis and presently invisible to Basenji.

### Disease-associated loci

Basenji’s utility for analyzing human genomic variation goes beyond intermediate molecular phenotypes like eQTLs to downstream ones like physical traits and disease. Basenji also offers substantial upside—eQTL analysis is highly informative for disease variant interpretation, but few cell types can plausibly be sampled to conduct such an investigation. With Basenji, a single experiment is sufficient to predict a genomic variant’s influence on gene expression in that cell type. We hypothesized that a predictive view of the 973 human samples profiled by CAGE would offer a novel perspective on disease variants.

To test the utility of Basenji SNP scores for this application, we acquired a curated set of disease variants studied by the successful DeepSEA method to predict variant influence on TF binding and chromatin (**Zhou and Troyanskaya 2015**). DeepSEA trained deep convolutional neural networks to predict ENCODE and Roadmap DNase-seq and ChIP-seq peak calls. The set includes 12,296 bi-allelic SNPs taken from the NIH GWAS Catalog database (**MacArthur et al. 2017**) and a negative set with matched minor allele frequencies that we sampled down to the same size. Because this experiment is cell/tissue-agnostic, we computed the penultimate layer predictions for each SNP allele in the 128 bp bins across the surrounding region. We assigned the SNP the maximum log_2_ ratio between allele values for each penultimate layer unit. 128 principal components were sufficient to represent the full profile well (Supplemental Fig S9). We maintained the same 10 cross validation folds reported by DeepSEA to compare methods. A logistic regression model to predict GWAS catalog presence using these Basenji principal component features achieved 0.666 AUROC, slightly greater than the .657 achieved by DeepSEA using a more complicated regression model that also included conservation statistics (p-value < 0.10 by dependent t-test over 10 folds) (Supplemental Fig S9). A joint model adding DeepSEA’s predictions as a feature to our logistic regression model increased AUROC to 0.706 (p-value < 1.6×10^−5^), supporting the orthogonal value added by Basenji SNP scores beyond the state of the art.

Having established meaningful signal in the predictions, we analyzed a set of 1170 loci associated with 40 autoimmune diseases and blood cell traits that were fine mapped using the linkage-based statistical method PICS (Farh et al. 2015). 67 loci contained a variant predicted to alter a gene’s transcription in one of the CAGE experiments by >10%, and an additional 73 contained a variant predicted to alter one of the chromatin profiles >10% at a gene’s start sites. rs78461372, associated with multiple sclerosis via linkage with the lead variant rs74796499 (**Lambert et al. 2013**), emerged from this analysis. Basenji predicts the C>G at rs78461372 to increase transcription of the nearby *GPR65* in many cells, most severely acute lymphoblastic leukemia cell lines, thyroid, insular cortex, and a variety of immune cells. *GPR65* is a receptor for the glycosphingolipid psychosine and may have a role in activation-induced cell death and differentiation of T-cells (**Wang et al. 2004**). Despite the fact that the multiple sclerosis GWAS literature does not highlight *GPR65*, sphingolipid metabolism has emerged as a therapeutic target for MS via the drug fingolimod, a sphingosine analogue that alters immune cell trafficking and is now in clinical use (**Brinkmann et al. 2010**). The model also predicts a small expression increase for *GALC*, a nearby gene whose start site is 12.9 kb away from the variant, in many immune cells. Both genes have been implicated in several immune diseases (inflammatory bowel disease, Crohn’s disease, ulcerative colitis) via variants independent of this set (**Koscielny et al. 2017**), and both genes may propagate a downstream causal influence on the disease.

PICS fine mapping assigns rs78461372 a 5% probability of causal association with multiple sclerosis and the lead variant rs74796499 a 24% probability. Basenji predicts no effect for rs74796499 or any other variants in the PICS credible set. To validate the predicted stronger effect of rs78461372 on nearby transcription, we consulted the GTEx eQTL analysis (The GTEx Consortium 2017). GTEx supports Basenji’s diagnosis, detecting significantly increased *GPR65* expression for individuals with the minor allele at rs78461372 in transformed fibroblasts (marginal beta 0.75; p-value 1.8×10^−9^); the competing correlated variant rs74796499 has a smaller measured effect (marginal beta 0.66; p-value 4.9×10^−7^).

To better understand the model prediction, we performed an in silico saturation mutagenesis (**Kelley et al. 2016; Lee et al. 2015**) in several affected cell types. That is, we generated sequences that introduce every possible mutation at all sites in the region, predicted CAGE activity, and computed the difference from the reference prediction. The functional motifs that drive the model’s prediction emerge as consecutive sites where mutations result in large differences. rs78461372 overlaps an ETS factor motif adjacent to a RUNX factor motif, where the G allele confers a stronger hit to the JASPAR database PWM for ETS, discovered using the motif search tool Tomtom (**Gupta et al. 2007; Mathelier et al. 2016**) (Figure 6). Interestingly, Basenji predicted opposite effects for these motifs in different cell types. In immune cells, where *GPR65* and *GALC* are more active, disruption of the motifs decreases the prediction. Alternatively, disruption of the same motifs increases the model’s prediction for expression in several tissues, including insular cortex. Altogether, Basenji predictions shed substantial light on this complex and influential locus, offering several promising directions for future work to unravel the causal mechanism.

**Figure 6.**
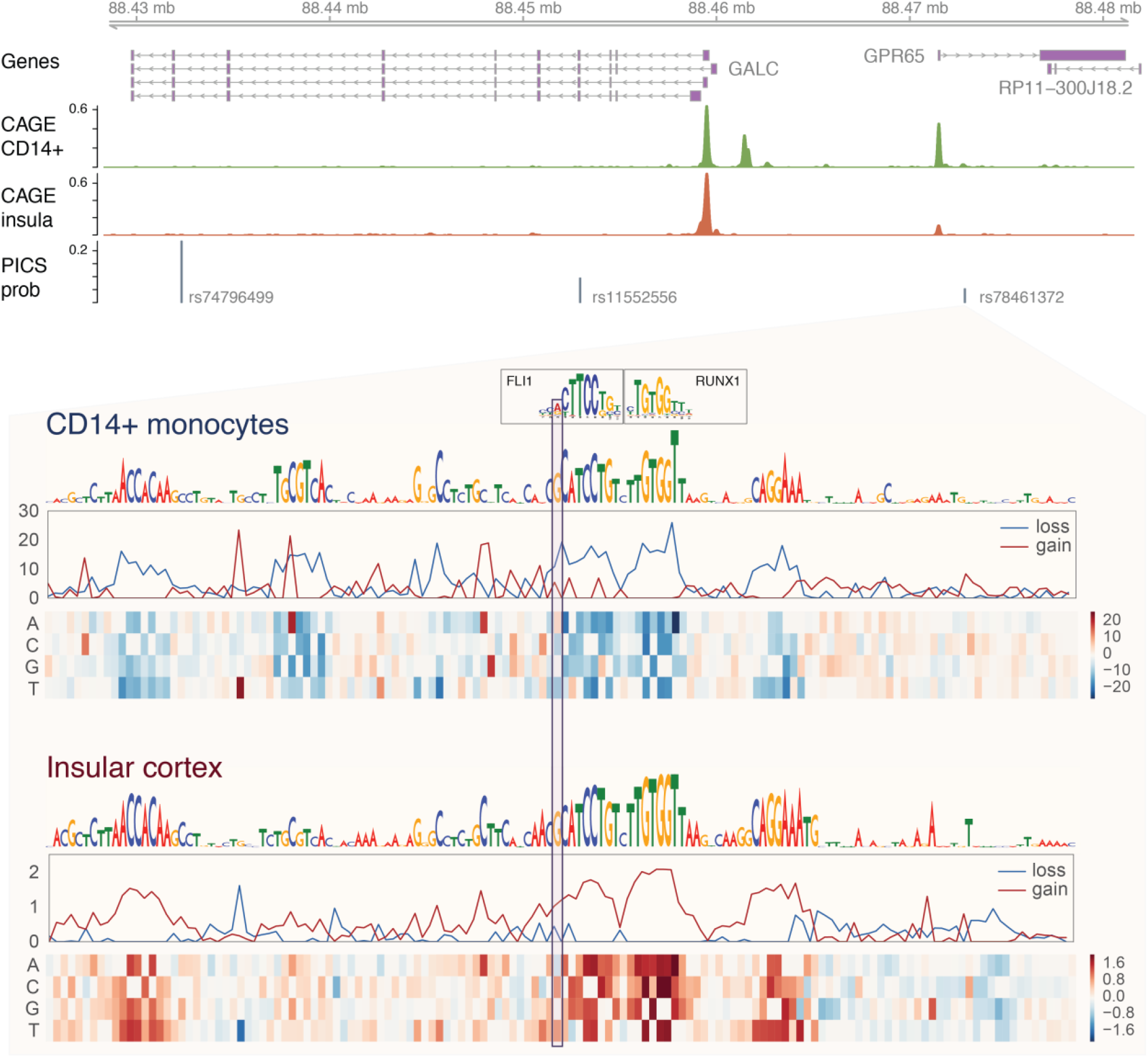
Basenji gene-specific variant scores illuminate multiple sclerosis associated locus. Lead variant rs74796499 is associated with multiple sclerosis (IMSGC 2013). Among the credible set of linked variants, Basenji predicts that rs78461372 would alter transcription of the nearby genes GPR65 and GALC. In immune cells, such as treated CD14^+^ cells depicted here, both genes are transcribed and the C>G introduces an ETS factor motif that enhances transcription. In contrast, in other cell types, e.g. insular cortex, where GPR65 is far less transcribed, Basenji predicts the same motifs play a role in repressing the gene.

## Discussion

Transcriptional regulation is the primary driver of gene expression specificity across cell types and states (**Waszak et al. 2015; Grubert et al. 2015; Battle et al. 2015; Li et al. 2016**). The genome research community needs more effective models of how sequence determines transcription in large mammalian genomes in order to understand how genomic alterations influence the downstream physical output of those genomes. Here, we introduced a comprehensive model to predict epigenetic and transcriptional profiles from DNA sequence. A deep convolutional neural network, trained on >4000 datasets, shares information across large distances with dilated layers to make sequential predictions along the chromosomes. The model explains considerable variance in these data, including cell type-specific activity. Predictions for sequences containing different versions of variant alleles statistically align with measurements made in human populations and subjected to eQTL analysis.

Although we demonstrated the present utility of this approach, there are several indications that we may be scratching the surface of what will be possible in this space. The datasets analyzed vary in quality, both by signal-to-noise ratio and technical variance from under-sampling with limited sequencing. We observed increasing predictive accuracy for experiments with greater sequencing depth and greater consistency between replicates. Thus, sequence-based modeling will benefit from improved experimental protocols, which are an area of active research (e.g., CUT&RUN in place of ChIP (**Skene and Henikoff 2017**)). Furthermore, most of the samples either describe cell lines or heterogeneous tissue samples. Pending efforts to profile more pure, specific cell types and states will enhance our ability to thoroughly detect all regulatory elements and offer precise predictions of when and where regulatory activity will occur (**Regev et al. 2017**). As the number of potential datasets grows, our simple multi-task training scheme that treats every (experiment, tissue/cell type, replicate) as an independent prediction task may not be optimal; more structured multi-task approaches that share information across dataset features offer a potential route to improvement (**Kang et al. 2011**). Finally, our training scheme involves a simplifying assumption that the reference genome specifies the DNA underlying the functional annotations studied. While experiment-specific genomes would provide more precise training data, these are mostly unavailable and would be very challenging computationally to incorporate. Advances that enabled such genome-specific training would likely improve the model.

Dilated convolutions extended the reach of our model to view distal regulatory elements at distances beyond previous models, achieving a 32 kb receptive field width. This distance contains many, but certainly not all regulatory elements (**Mifsud et al. 2015; Javierre et al. 2016**). Thus, extending the model’s vision and exploring architectures to better capture the long-range logic of gene regulation are a promising avenue for future research, with considerable potential to improve predictive accuracy and interpret variants far from genes.

Despite focusing only on transcription without considering post-transcriptional regulation and interaction across regulatory layers (**Skalska et al. 2017**), we found our predictions highly informative of which genomic variants would be highlighted as eQTLs in population studies measuring RNA abundance levels. We foresee considerable potential in further integrating regulatory activity models trained on functional genomics profiles with population genotyping and phenotyping. These orthogonal approaches both offer views into how the noncoding genome works, and their joint consideration ought to sharpen those views. We envision Basenji as an important step forward in this direction.

## Methods

### Data preprocessing

Finer resolution analysis of large genomic regions exposes expressive machine learning models more to biases in functional genomics sequencing experiments (e.g. fragment GC%) (**Meyer and Liu 2014; Benjamini and Speed 2012**) and repetitive DNA. Processed data available for download by the consortiums disposes multi-mapping reads and largely ignores these biases. Thus, we carried out our own processing of this data, with greater care taken to account for how these factors would influence the downstream training algorithms.

We downloaded FASTQ files for 973 CAGE experiments performed by FANTOM5 (**Forrest et al. 2014**), 593 DNase-seq and 1704 histone modification ChIP-seq performed by ENCODE (**The ENCODE Project Consortium et al. 2012**), and 356 DNase-seq and 603 histone modification ChIP-seq performed by the Epigenomics Roadmap (**Roadmap Epigenomics Consortium et al. 2015**) (Supplemental Table 1). We aligned the reads with Bowtie 2, requesting the program return a maximum of 10 multi-mapping alignments (**Langmead and Salzberg 2012**). We proportioned these multi-reads among those 10 positions using an EM algorithm that leverages an assumption that coverage will vary smoothly (**Zhang and Keleş 2014**). In the algorithm, we alternate between estimating expected coverage across the genome using a Gaussian filter with standard deviation 32 and re-allocating multi-read weight proportionally to those coverage estimates. We normalized for GC% bias using a procedure that incorporates several established ideas (**Benjamini and Speed 2012; Teng and Irizarry 2016**). We assigned each position an estimated relevant GC% value using a Gaussian filter (to assign greater weight to nearby nucleotides more likely to have been part of a fragment relevant to that genomic position). Then we fit a third-degree polynomial regression to the log_2_ coverage estimates. Finally, we reconfigured the coverage estimates to highlight the residual coverage unexplained by the GC% model. A Python script implementing these procedures to transform a BAM file of alignments to a BigWig file of inferred coverage values is available in the Basenji tool suite.

Avoiding assembly gaps and unmappable regions >1 kb, we extracted (2^17^=) 131 kb nonoverlapping sequences across the chromosomes. We discarded sequences with >35% unmappable sequence, leaving 14,533 sequences. We randomly separated 5% for a validation set, 5% for a test set, and the remaining 90% for training. Within each sequence, we summed coverage estimates in 128 bins to serve as the signal for the model to predict.

### Model architecture and training

We implemented a deep convolutional neural network to predict the experimental coverage values as a function of the underlying DNA sequence. The high-level structure of the network consisted of convolution layers, followed by dilated convolution layers, and a final convolution layer (Figure 1). All layers applied batch normalization, rectified linear units, and dropout. Standard convolution layers applied max pooling in windows of 2, 4, 4, and 4 to reach the 128 bp bin size. We compared the predicted and measured values via a Poisson regression log-likelihood function. We used Tensorflow implementations for these layers (**Abadi et al. 2016**).

We chose 128 bp because it corresponds to the power of two closest to the 146 bp distance between nucleosome core particles. It offers a finer resolution view than the popular ChromHMM method, which studies these data at 200 bp resolution (**Ernst and Kellis 2010**). Greater sequencing depth for the training datasets would enable enhanced resolution in future studies.

Dilated convolutions are convolution filters with gaps whose size increases by a factor of two in each layer, enabling the receptive field width to increase exponentially (**Yu and Koltun 2015**). Dense connection of these layers means that each layer takes all previous layers as input, as opposed to taking only the preceding layer (**Huang et al. 2016**). This architecture allows for far fewer filters per layer because the incoming representation from the standard convolutional module and the subsequent refinements of the dilated layers are all passed on; this allows each layer to focus on modeling the residual variance not yet captured (**He et al. 2015**). We applied seven dilated convolution layers in order to reach a receptive field width of ~32 kb. This width will capture only a subset of possible distal regulatory interactions (**Mifsud et al. 2015; Javierre et al. 2016**), and we intend to engineer methods to increase it. Nevertheless, it captures substantially more relevant regulatory sequence than previous models, and evidence that variant effect magnitude decreases with distance suggests there will be diminishing returns to extension (**The GTEx Consortium 2017**).

We initialized weight values using Glorot initialization (Glorot and Bengio) and optimized the loss function via stochastic gradient descent with learning rates adapted via ADAM (**Kingma and Ba 2014**). Our TensorFlow implementation leverages automatic differentiation and the chain rule to compute the gradient of the loss function with respect to each parameter to step towards a local optimum (**Abadi et al. 2016**). We used Bayesian optimization via the GPyOpt (https://github.com/SheffieldML/GPyOpt) package to search for effective hyper-parameters throughout the model, including the convolution widths, convolution filter numbers, dropout rates, learning rate, and momentum parameters (**Snoek et al. 2012**).

During training, we applied two strategies to augment the data and reduce overfitting. Every other epoch, we reverse complemented the DNA sequences and reversed the values. We also iterated over minor shifts of 0, 1, 2, and 3 nucleotides left and right each epoch. During testing, we averaged predictions over these transformations.

### Peaks binary classification comparison

To benchmark Basenji versus our previous approach Basset, we transformed the training and test datasets to binary peak calls on shorter sequences and trained a Basset model. We divided each 131 kb sequence into 1024 bp subsequences. For each dataset, we called peaks on the smoothed, normalized counts in the center 256 bp of the subsequences using a Poisson model parameterized by the maximum of a global and local null lambda similar to the MACS2 approach and applied a 0.01 FDR cutoff (**Zhang et al. 2008**). We trained the Basset model using the most effective hyper-parameters yet discovered, which were used in a recent application (**Reshef et al. 2017**).

### Gene expression cluster comparison

To measure Basenji’s ability to recapitulate gene expression clusters from the experimental data, we focused on the 2000 most variable genes and sampled sets of 1000 using a bootstrap procedure. For each sample, we performed Gaussian mixture model clustering with 10 clusters on the experimental and predicted gene expression matrixes across cell types. We quantified the similarity of the clusters with the adjusted Rand index statistic. The distribution of the statistic was approximately normal; thus, we estimated the mean and variance of the distribution to compute a p-value that the distribution was greater than zero.

### Regulatory element saliency maps

We desired a computationally efficient measurement of the influence of distal sequence on gene expression predictions, focusing on the 128 bp resolution representations after the convolution and pooling layers, but before the dilated convolutions. Deep learning research has suggested several effective schemes for extracting this information. Guided by the insights of prior work (**Shrikumar et al. 2017; Bach et al. 2015**), we computed experiment-specific saliency maps as the dot product of the 128 bp bin representations and the gradient of the model prediction summed across the sequence with respect to those bin representations. The rectified linear unit nonlinearity guarantees that all representation values will be positive. Thus, positive gradients indicate that stronger recognition of whatever triggered the unit would increase the prediction (and weaker recognition would decrease it); negative gradients indicate the opposite. Taking the dot product with the sequence’s representation amplifies the signal and sums across the vector, aggregating the effect into one signed value. Positive values identify regions where activating elements were recognized, and negative values identify repressor elements.

We applied the following procedure to compare saliency scores to experimental regulatory element annotations. We downloaded ENCODE annotations for GM128768 promoter (ENCFF492VIP), enhancer (ENCFF811LAE), and CTCF (ENCFF002COQ) sites as BED files. We subtracted the promoters from the enhancers, and we subtracted both the promoters and enhancers from the CTCF sites. We formed a background set by shuffling these annotations within the 131 kb regions for which we computed scores and removing instances that overlapped true elements. We computed a p-value for every saliency score as the probability that a normal distribution fit to the background scores would have a more extreme value and corrected for multiple hypotheses with the Benjamini-Hochberg false discovery rate procedure to 0.05.

### GTEX eQTL analysis

We downloaded the eQTL analysis in the GTEx V6p release and primarily studied the chi-squared statistics and significance calls (**The GTEx Consortium 2017**). Nearby variants in the population data can have highly correlated statistics due to linkage disequilibrium. In contrast, Basenji can isolate the contribution of individual variants. To place SED scores and eQTL statistics on a level plane, we computed their signed LD profile (**Reshef et al. 2017**). The signed LD profile of a signed genomic annotation gives the expected marginal correlation of each SNP to a hypothetical phenotype for which the true causal effect of each SNP is the value of the annotation at that SNP. It is what we would expect the GWAS summary statistics to look like it Basenji predictions were exactly correct for each gene. We computed the signed LD profile by multiplying the SED scores by a population LD matrix estimated from the 1000 Genomes Phase 3 Europeans (**The 1000 Genomes Project Consortium 2015**).

We included several covariates that are known to influence eQTL chi-squared statistics—LD score and TSS distance (**Liu et al. 2017**). LD score measures the amount of variation tagged by an individual variant (**Bulik-Sullivan et al. 2015**). We found that LD score correlated with the chi-squared statistics. Thus, we downloaded pre-computed scores for the European 1000 Genomes from the LDSC package (**Bulik-Sullivan et al. 2015**) and included them in the analyses. We also found that distance to the nearest TSS correlated with the chi-squared statistics; variants closer to TSSs are more likely to influence gene expression. To control for this effect, we annotated SNPs with indicator variables classifying TSS distance as < 500 bp, 500-2000 bp, 2000-8000 bp, or 8000-32000 bp and computed LD scores to each annotation using LD information from 1000 Genome Phase 3 Europeans, as in (**Finucane et al. 2015**).

Finally, we pruned the set of variants down to ensure that there were no pairs of variants in strong LD (R^2^ > 0.5). In a first analysis, we fit regression models individually for each tissue with LD score and |SED-LD| to the chi-squared statistics, and considered the significance of coefficients assigned to |SED-LD| across all variants. In a second analysis, we added the TSS-LD variables to the regression.

## Software availability

The code to preprocess data, train models, and perform the analyses described is available in the Supplemental Material and from https://www.github.com/calico/basenji.

## Acknowledgments

We thank David Hendrickson, Daphne Koller, Geoffrey Fudenberg, and Marta Mele for feedback on the manuscript. We thank Ben Passarelli for invaluable computing hardware support.

## Competing financial interests

DRK is employed by Calico LLC. DB, CYM, JS, and MB are employed by Google Inc.

