## Supplementary Materials for "Sequential regulatory activity prediction across chromosomes with convolutional neural networks"

### Title

1. Calico Labs. South San Francisco, CA, USA.
2. Department of Computer Science. Harvard University. Cambridge, MA, USA.
3. Google Brain. Cambridge, MA, USA.

### Correspondence

David R. Kelley

1170 Veterans Blvd.

South San Francisco, CA 94080



### Keywords

Machine learning, Convolutional neural network, Gene expression, Gene regulation, Functional genomics

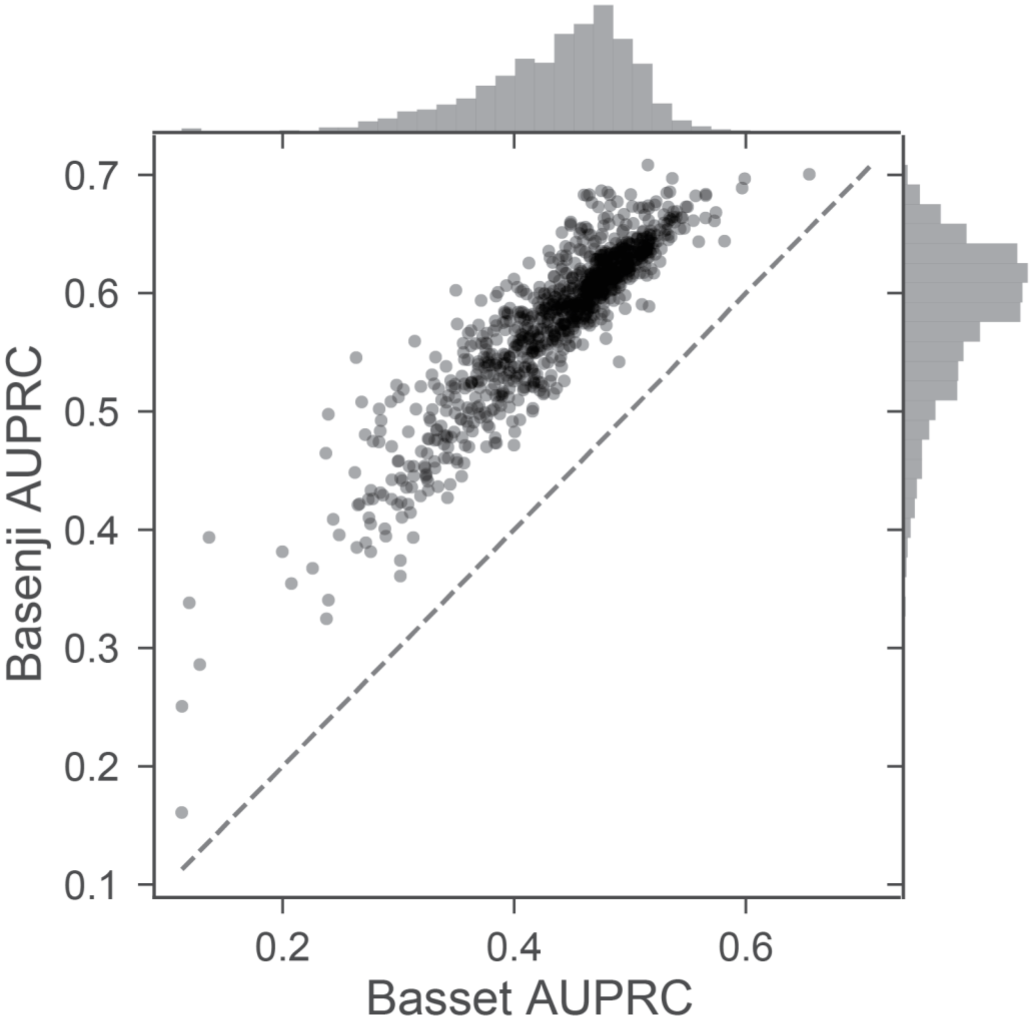


*Supplementary Figure S1 - Basenji increases peak call accuracy over Basset.*

We trained a Basset model to predict peak calls in 949 DNase-seq experiments (Methods). For Basset and Basenji, we plotted the area under the precision-recall curve (AUPRC) for each experiment's predictions. The advances introduced in the Basenji model increase the average AUPRC from 0.435 to 0.577 and median from 0.449 to 0.591.

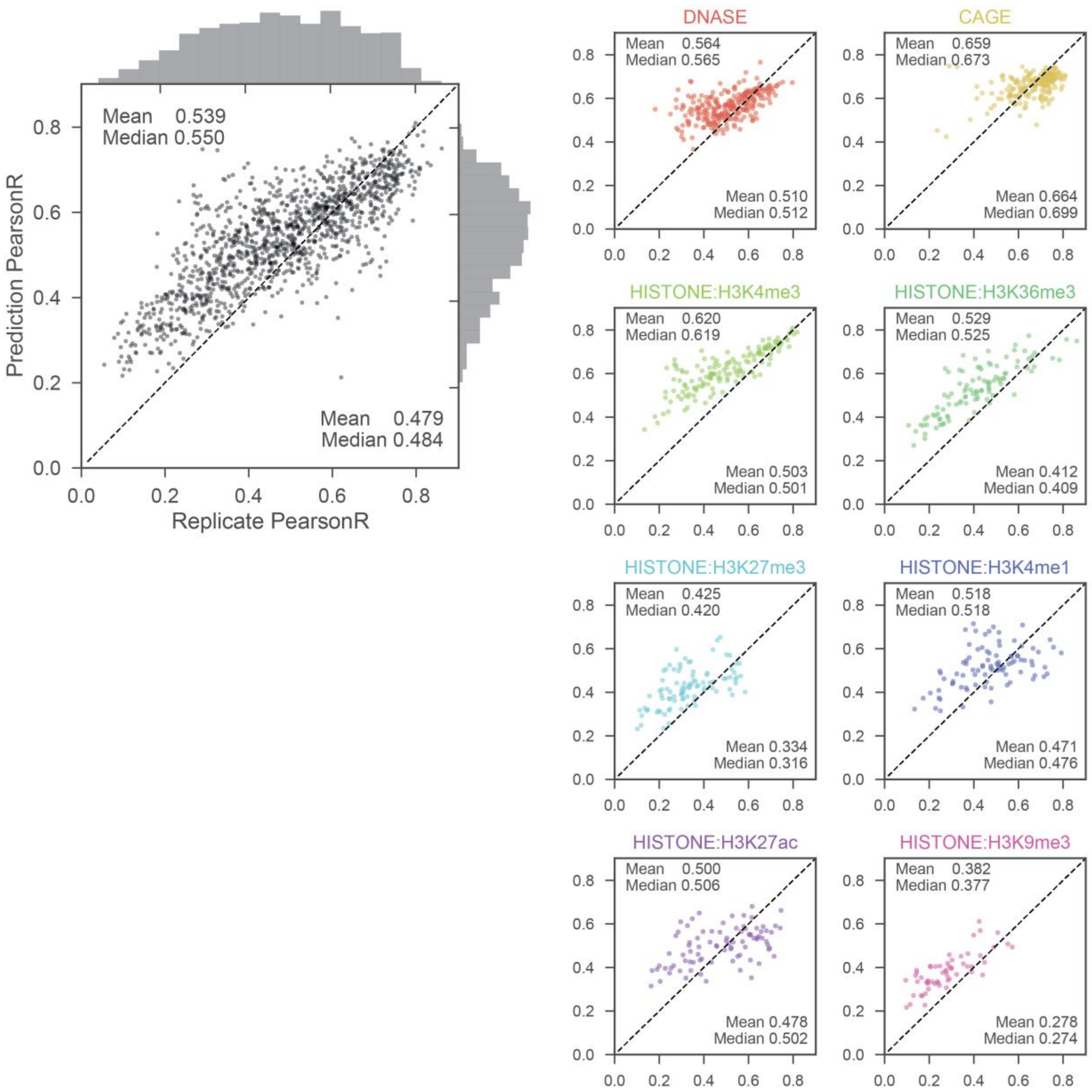


*Supplementary Figure S2 – Basenji predictions within-replicate match replicate concordance.* For all replicated experiments, we plotted log-log Pearson correlation between the replicate experiments versus the correlation between the experiment and prediction (averaged across replicates). On the right, we make the same plots, faceted by experiment type.

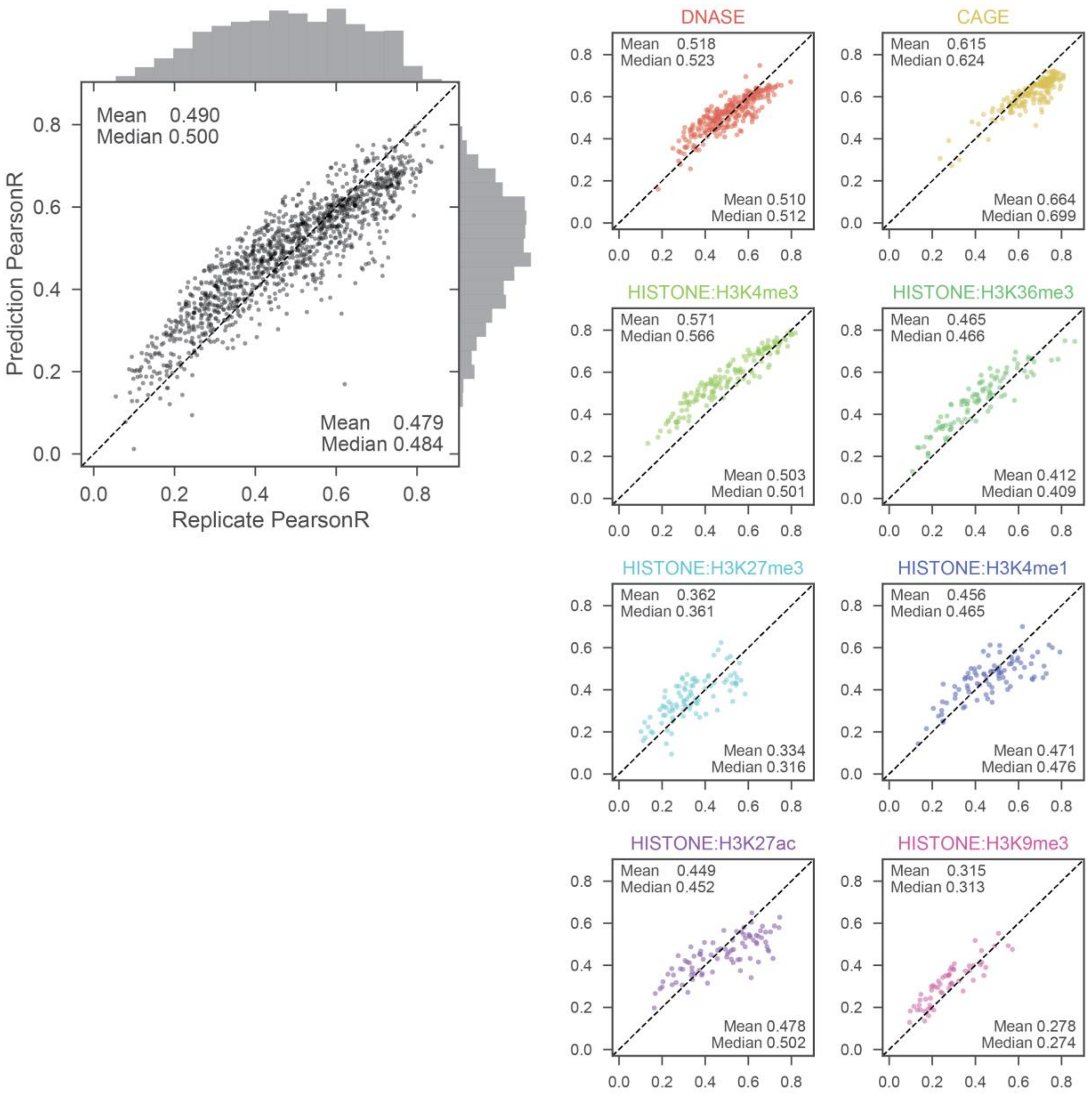


*Supplementary Figure S3 – Basenji predictions cross-replicate match replicate concordance.* For all replicated experiments, we plotted log-log Pearson correlation between the replicate experiments versus the correlation between the experiment and its replicate's prediction (averaged across replicates). On the right, we make the same plots, faceted by experiment type.

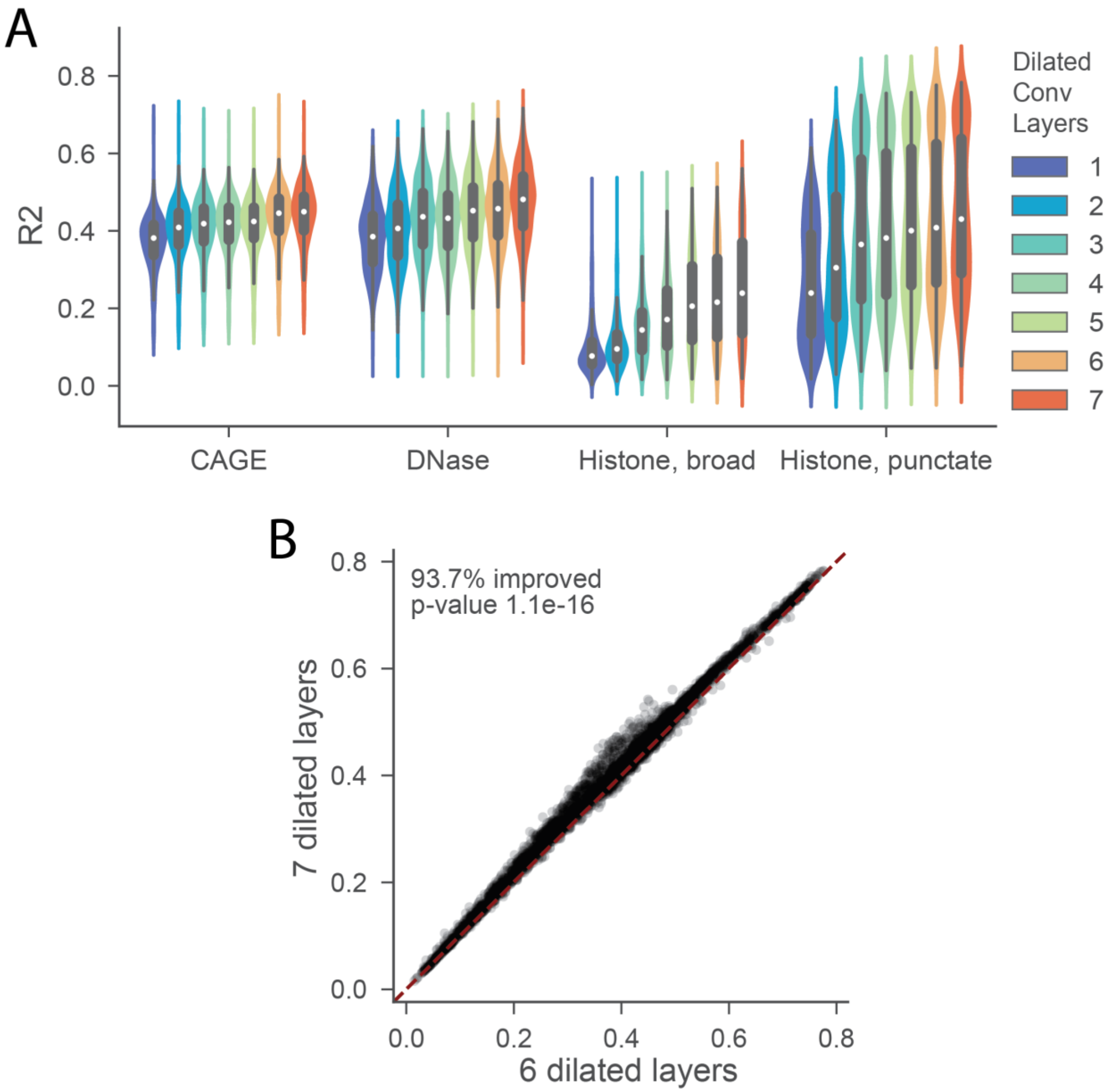


*Supplementary Figure S4 - Dilated layers improve predictive accuracy.*

We trained models for a range of dilated convolution layer number. (A) We plotted the distribution of test R2 for each experiment, by data type. Test accuracy increases with each additional layer for all data types. (B) We plotted the test R2 of each experiment for the 6 layer versus 7 layer model. Adding the 7$^{th}$ layer improves test accuracy for 93.7% of the datasets. An 8$^{th}$ layer would reach outside the bounds of the sequence too frequently for this input length, but may add value for larger sequences.

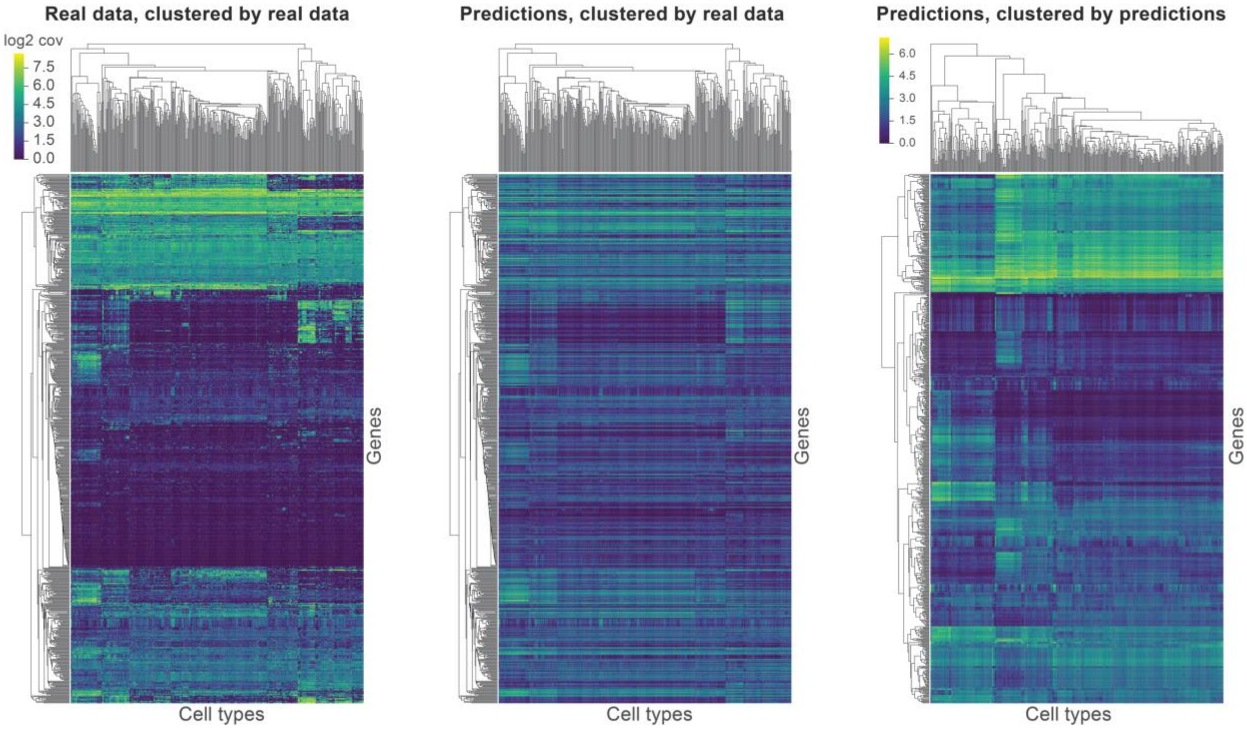


*Supplementary Figure S5 - Predictions maintain cell type-specific expression clusters.*

On the far left, we performed hierarchical clustering and plotted as a heat map the experimental gene expression matrix across cell types after quantile normalization. On the far right, we similarly plotted the Basenji gene predictions matrix. In the center, we froze the row and column order from the experimental data clustering on the left and substituted in Basenji predictions. Although the sharp definitions smear, the clusters remain visible. We used Euclidean distance and average linkage in the hierarchical clustering.

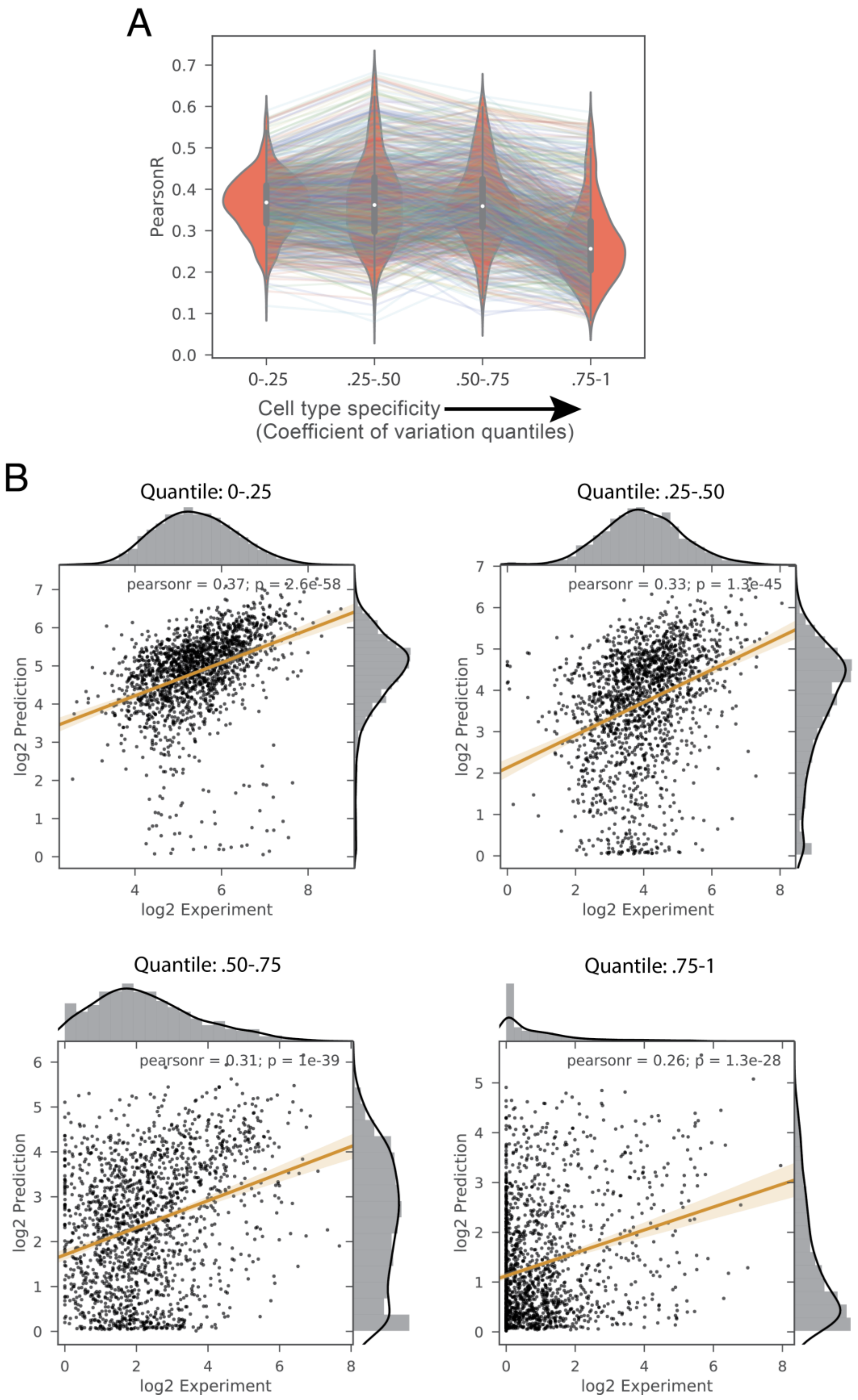
A
PearsonR
0.0
0.1
0.2
0.3
0.4
0.5
0.6
0.7
0-.25
.25-.50
.50-.75
.75-1
Cell type specificity
(Coefficient of variation quantiles)
B
Quantile: 0-.25
pearsonr = 0.37; p = 2.6e-58
log2 Prediction
log2 Experiment
Quantile: .25-.50
pearsonr = 0.33; p = 1.3e-45
log2 Prediction
log2 Experiment
Quantile: .50-.75
pearsonr = 0.31; p = 1e-39
log2 Prediction
log2 Experiment
Quantile: .75-1
pearsonr = 0.26; p = 1.3e-28
log2 Prediction
log2 Experiment

*Supplementary Figure S6 – Cell type-specific gene accuracy.*

We established a set of 5198 with sufficient expression to compute meaningful variance statistics as those with mean > 0.5 and max > 3 expression across CAGE samples after log transform and quantile normalization. We ranked these genes by their coefficient of variation across samples and formed four quantile sets containing ~1300 genes each. For each CAGE sample, we computed the Pearson correlation between the normalized experimental measurements and Basenji predictions within each quantile set. (A) We plotted each CAGE sample's quantile correlations as a line, and used violin plots to represent the four distributions. Correlation is stable until the most variable gene set, where the mean correlation decreases from 0.3673 (median 0.3592) in the third quantile to 0.2708 (median 0.2562) in the fourth. (B) We ranked the CAGE samples by the ratio of this decrease and chose the median sample to scatter plot the genes' normalized experimental measurements and predictions in each quantile.

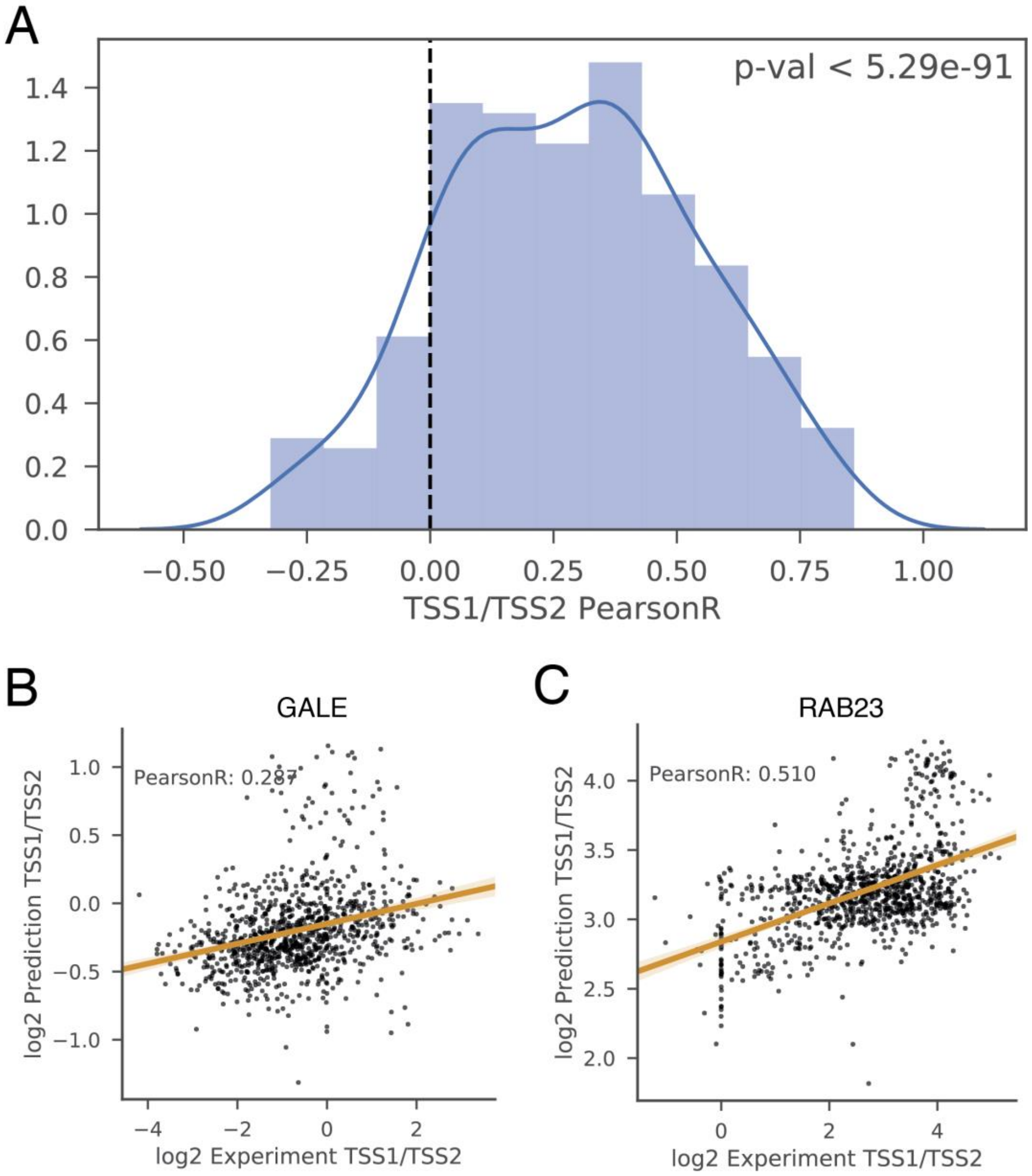


*Supplementary Figure S7 – Predictions reflect alternative TSS usage.*

We delineated a set of 201 genes with multiple distinct TSSs >500 bp, each with variance across CAGE samples >1 (for log-transformed, quantile normalized values). For each gene, we computed the log ratio of the most variable TSS's activity versus the second most variable TSS's activity. Then, we computed the Pearson correlation of this statistic for the experimental measurements and Basenji predictions across all CAGE samples. (A) We plotted a histogram of these correlations. The set has greater mean than zero by T-test with p-value $<1 \times 10^{-90}$. (B) We plotted the log ratio of TSS1/TSS2 activity for the experimental measurement versus prediction for the median correlation gene *GALE* (C) and the 75$^{th}$ percentile gene *RAB23*.

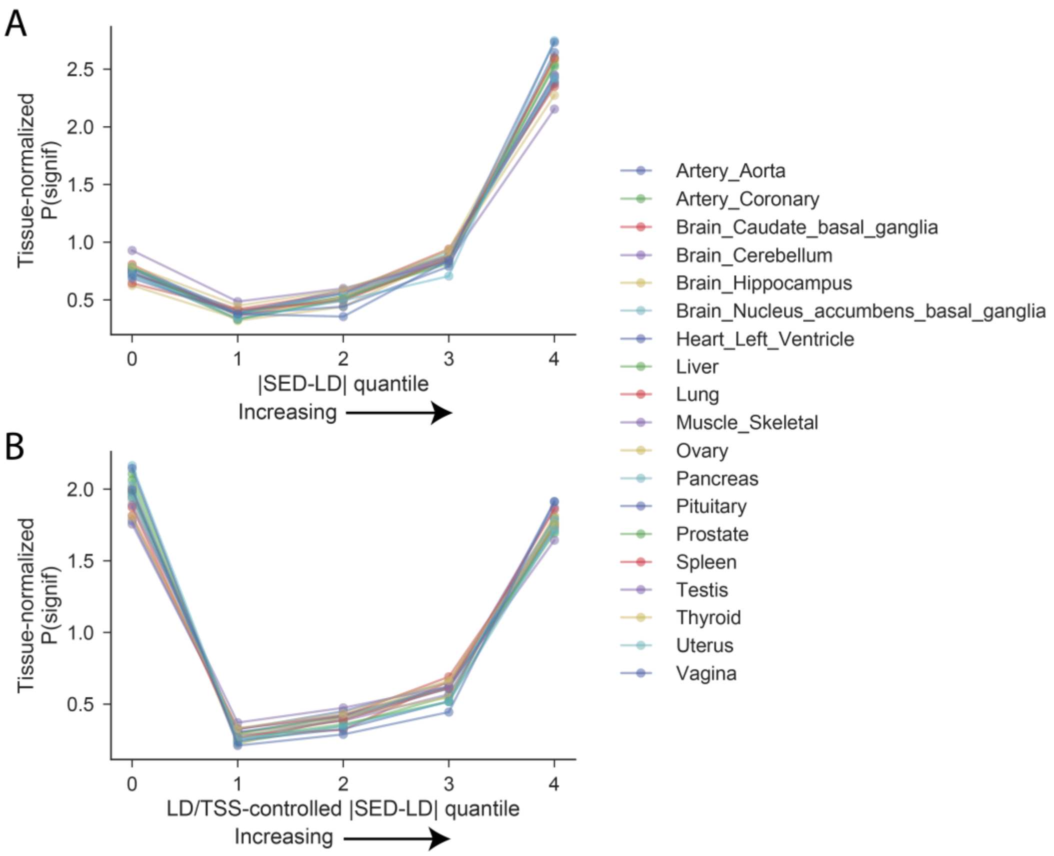


*Supplementary Figure S8 – SNP expression difference predictions relate to GTEx eQTL statistics.* We distributed SED by the LD correlation matrix to more readily compare to eQTL measurements in human populations (Methods). |SED-LD| shows a strong relationship with eQTL statistics from GTEx. (A) For each tissue, we ranked the variants by the difference between their regression predictions including and excluding |SED-LD| and formed five quantiles. We computed the proportion of significant eQTLs in each quantile and divided by the proportion of all variants called eQTLs in that tissue to normalize the tissues to a level plane. The line plots show those normalized significance proportions in each quantile, which rise to 3.2-5.8x over the average of the bottom three quantiles in all 19 tissues. (B) We observed that TSS distance also related to variant eQTL statistics and recomputed the regression-based ranking and quantiles including TSS distance covariates (Methods). The highest SED-LD quantile remains highly enriched for eQTLs. Enrichment of the lowest quantile may be attributable to variants that influence gene expression via mechanisms beyond the transcriptional regulation that Basenji focuses on (GTEx Consortium 2017; Battle et al. 2015). Variants that affect post-transcriptional mechanisms such as splicing would collect in the lowest quantile where the SNPs tag substantial variation near the gene, but have low |SED-LD| predictions.

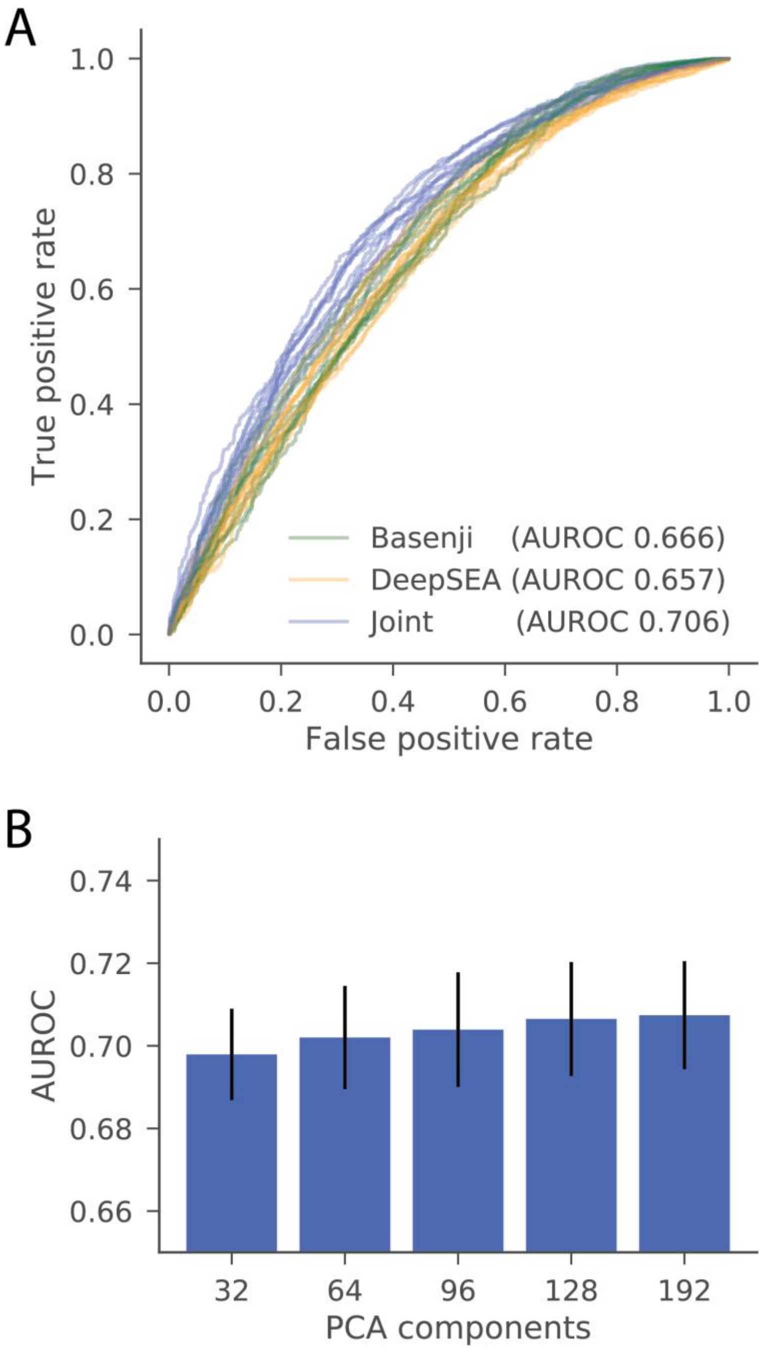


*Supplementary Figure S9 – Basenji predictions exceed previous methods for GWAS classification.* We computed Basenji SNP scores for a dataset containing 12,296 bi-allelic SNPs taken from the NIH GWAS Catalog database (MacArthur et al. 2017) and a negative set with matched minor allele frequency. We trained a logistic classifier to predict presence in the GWAS catalog. The Basenji model matches DeepSEA, whose authors included conservation statistic features, too. A joint model adding DeepSEA's predictions as a feature to ours achieves significantly greater accuracy than either alone
